# Description of a widespread bacterial secretion system with chemically diverse substrates

**DOI:** 10.1101/2020.01.20.912956

**Authors:** Alex S. Grossman, Terra J. Mauer, Katrina T. Forest, Heidi Goodrich-Blair

## Abstract

In host-associated bacteria, surface and secreted proteins mediate acquisition of nutrients, interactions with host cells, and specificity of tissue-localization. In Gram-negative bacteria, the mechanism by which many proteins cross or become tethered to the outer membrane remains unclear. The <u>d</u>omain of <u>u</u>nknown function (DUF)560 occurs in outer membrane proteins throughout Proteobacteria and has been implicated in host-bacteria interactions and lipoprotein surface exposure. We used sequence similarity networking to reveal three subfamilies of DUF560 homologs. One subfamily includes those DUF560 proteins experimentally characterized to date: NilB, a host-range determinant of the nematode-mutualist *Xenorhabdus nematophila*, and the <u>s</u>urface lipoprotein <u>a</u>ssembly <u>m</u>odulators Slam1 and Slam2, which facilitate msurface exposure of lipoproteins in *Neisseria meningitidis* (1, 2). We show that DUF560 proteins from a second subfamily facilitate secretion of soluble, non-lipidated proteins across the outer membrane. Using *in silico* analysis, we demonstrate that DUF560 gene complement correlates with bacterial environment at a macro level and host association at a species level. The DUF560 protein superfamily represents a newly characterized Gram-negative secretion system capable of lipoprotein surface exposure and soluble protein secretion with conserved roles in facilitating symbiosis. In light of these data, we propose that it be titled the type <u>eleven s</u>ecretion <u>s</u>ystem (TXISS).

**Importance:** The microbial constituents of a host associated microbiome are decided by a complex interplay of microbial colonization factors, host surface conditions, and host immunological responses. Filling such niches requires bacteria to encode an arsenal of surface and secreted proteins to effectively interact with the host and co-occurring microbes. Bioinformatic predictions of the localization and function of putative bacterial colonization factors are essential for assessing the potential of bacteria to engage in pathogenic, mutualistic, or commensal activities. This study uses publicly available genome sequence data, alongside experimental results from representative gene products from *Xenorhabdus nematophila*, to demonstrate a role for DUF560 family proteins in the secretion of bacterial effectors of host interactions. Our research delineates a broadly distributed family of proteins and enables more accurate predictions of the localization of colonization factors throughout Proteobacteria.

## Introduction

All plants and animals exist in association with bacterial symbionts that can contribute to nutrition, protection, development, and reproduction. These symbionts express surface and secreted proteins that facilitate host interactions through a variety of functions, including acquisition of nutrients (e.g. TonB-dependent transporters in Gram-negative host-associated bacteria (3, 4)), interaction with host cells (e.g. *Acinetobacter baumanii* OmpA binding host desmoplakin (5)), and specificity in host-range and tissue-localization (e.g. *Vibrio fischeri* pilin variability mediating host specificity (6)).

Recently, a mechanism of lipoprotein surface tethering was identified in the human pathogen *Neisseria meningitidis* and termed the Slam (<u>S</u>urface lipoprotein <u>a</u>ssembly <u>m</u>odulator) machinery (1, 2). Slam proteins containing the β-barrel domain DUF560 are required for surface presentation of certain lipoproteins, including those that capture metal-carrying compounds used by hosts to sequester nutrients from bacteria (1). Two *N. meningitidis* Slam proteins have been characterized, each with distinct lipoprotein substrates. Slam activity also has been demonstrated for DUF560 representatives from pathogens *Pasteurella multocida, Moraxella catarrhalis*, and *Haemophilus influenzae* (1, 2). However, most Slam homologs have no bioinformatically predicted substrate to date, and one study has found that *N. meningitidis* Slam1 can surface expose unlipidated fHbp variants (7). Thus, the full functional potential of DUF560 proteins is not yet known.

The DUF560 homolog NilB is a host-association and species-specificity factor in the nematode symbiont *Xenorhabdus nematophila*, a proteobacterium in the family *Morganellaceae* (8-10). A screen for *X. nematophila* mutants defective in colonizing *Steinernema carpocapsae* intestines revealed the <u>N</u>ematode intestinal localization locus (Nil)(10, 11). The Nil locus contains the genes *nilB* and *nilC*, each of which is independently necessary for colonization of nematodes. Biochemical and bioinformatic analyses have established that NilC is an outer-membrane-associated lipoprotein and NilB is an outer-membrane β–barrel protein in the DUF560 protein family with a ∼140 amino acid periplasmic N-terminal domain that contains tetratricopeptide repeats (11-14).

To begin to understand the range of functions of DUF560 proteins, we assessed their ecological distribution, genomic context, and relatedness. We experimentally examined the *X. nematophila* DUF560 homolog HrpB, which is not predicted to interact with a lipoprotein substrate. Finally, to better understand the potential role of DUF560 proteins in host-symbiont interactions we analyzed DUF560 distribution among symbiotic *Xenorhabdus*. Our data demonstrate that the activities of the DUF560 family extend beyond lipoprotein surface presentation and constitute a new type <u>XI</u> bacterial <u>s</u>ecretion <u>s</u>ystem (TXISS) which, like the type II secretion system, is capable of acting on either membrane anchored or soluble proteins (15).

## Results

### TXISS cluster according to environment

Using homology to NilB or Slam proteins, previous work identified a wide distribution of DUF560 proteins within mucosal associated bacteria (2, 10, 11). To quantifiably delineate subfamilies within the TXISS we generated a <u>s</u>equence <u>s</u>imilarity <u>n</u>etwork (SSN) using the Enzyme Function Initiative toolset (EFI) (16-18) and annotated it to highlight environmental source or taxonomic grouping of the isolates containing DUF560 homologs (Fig 1; Supplemental Table 1). In this network analysis, protein sequences with a high identity (40% or greater) were gathered into data points called nodes and connected together with edges based on sequence similarity. Using all homologs in Interpro 73 and Uniprot 2019-02, we identified 10 major clusters of TXISS proteins. Cluster 1 was chosen for in-depth analysis since it contained the majority of nodes in the network (62.4%) and could be visually divided into three subclusters (1A, 1B, and 1C) using force directed node placement (Fig. 2). The remaining clusters displayed a preponderance of water and soil associated organisms and contained no characterized proteins. Consistent with our previous observations (11), the majority of cluster 1 nodes (75%) comprise sequences from various animal-associated isolates, while another 20% contain sequences from marine, freshwater, soil, or built-environment isolates.

**Figure 1.**
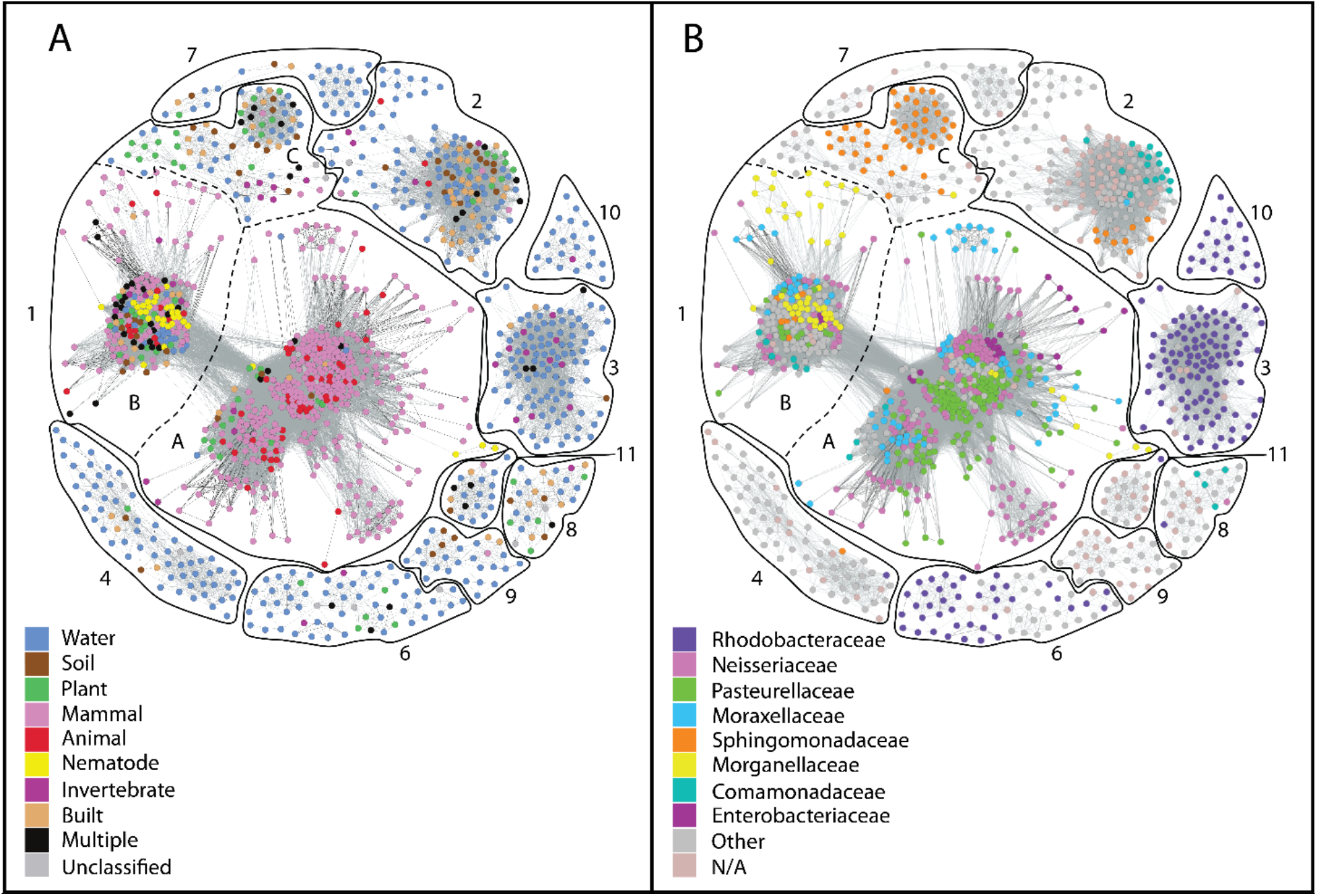
Sequence similarity network of all DUF560 proteins. SSN of DUF560 homologs generated by EFI-EST as accessed on 4.24.19 (16-18). Edges were drawn using an alignment score of 38, and any sequences which shared ≥ 40% identity were placed in a single node to allow the separation of clusters. Each node represents a group of highly similar sequences, with edge darkness demonstrating similarity, and the distance between nodes being determined via the Fruchterman-Reingold algorithm to optimize edge lengths (53). Each node was color coded to show the isolates’ environmental origin(s) (A) and taxonomic class (B).

**Figure 2.**
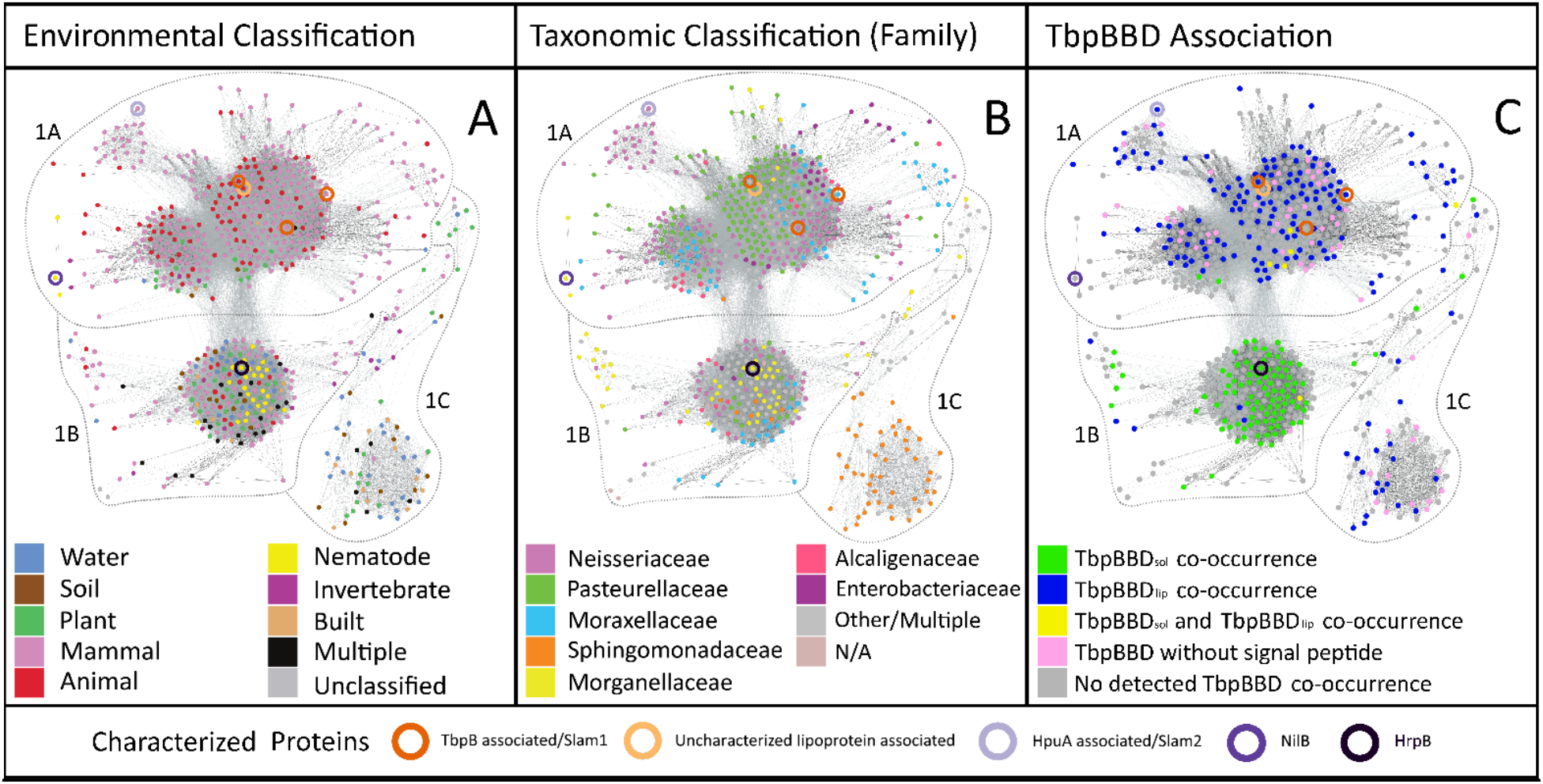
Cluster 1 of the TXISS sequence similarity network (SSN) forms subclusters according to environment of isolation and signal sequence of predicted cargo. All nodes from cluster 1 of the TXISS SSN were positioned using the Fruchterman-Reingold algorithm. The resulting graph was annotated according to sequences A) environmental origin(s) or B) taxonomic class of the isolates, or C) whether the node homolog(s) co-occur with a TbpBBD-domain encoding gene. Nodes containing experimentally characterized proteins are highlighted using colored circles as indicated at the bottom of the figure.

The division of nodes among the three subclusters more strongly reflected environmental origin than bacterial taxonomy (Fig. 2A). Subcluster 1A almost exclusively comprises animal-associated bacteria and contains all previously characterized TXISS, which separate according to predicted substrate when analyzed with higher stringency (Supplemental Fig. 1). Subclusters 1B and 1C have no previously characterized representatives. Subcluster 1B contains a mixture of sequences from host-associated and free-living bacteria, while subcluster 1C contains sequences largely from environmentally isolated *Sphingomonadaceae*. Subclusters 1A and 1B do not correlate well with taxonomy of the isolates (Fig. 2B). For example, cluster 1 contains 81 nodes with *Moraxellaceae* sequences. Of these, 79% are predominantly animal-associated and 12.3% are predominantly environmental isolates. The animal-associated isolates are enriched in subcluster 1A and environmental-isolates are enriched in subcluster 1B. These data demonstrate a correlation with lifestyle (e.g. free-living vs host associated) as opposed to taxonomy and suggest that subclusters indicate divergent molecular functions.

### TXISS cluster according to substrate

The Slam acronym was defined on the basis that DUF560 homologs from *N. meningitidis* facilitate surface exposure of lipoproteins, such as TbpB, LbpB, HpuA, and fHbp which are frequently encoded nearby (1, 2). The lipid tail is not essential for Slam-dependent surface exposure of a target (1, 7, 19). This result prompted us to consider whether co-occurrence with lipoproteins is a constant throughout cluster 1. We used the EFI Genome-Neighborhood Tool (16, 18, 20) to assay the genomic context of each subcluster. This analysis corroborated previous work demonstrating genomic association of DUF560 proteins with TonB, TonB-dependent receptors and proteins that have a Pfam TbpB_B_D domain, which will be referred to hereafter as TbpBBD (2, 21).

Given the prevalence of TbpBBD domains in the genome neighborhoods of DUF560 genes, and their known occurrence in lipoproteins surface exposed by Slams we examined their gene structures to detect potential patterns. Using a combination of genome-neighborhood-network data (18), Rapid ORF Description & Evaluation Online (RODEO) data (22), and manual annotation we analyzed all TbpBBD domain proteins co-inherited with sequences present in cluster 1 (Supplemental Table 2). The majority of TbpBBD-bearing proteins associated with subcluster 1A are predicted to be lipidated and have two TbpBBD domains similar to TbpB in *N. meningitidis* (2), therefore they will be referred to as TbpBBDlip. In contrast, the TbpBBD-bearing proteins associated with subcluster 1B are almost exclusively predicted to be soluble proteins and only have a single TbpBBD domain similar to hemophilin in *Haemophilus haemolyticus*, therefore they will be referred to as TbpBBDsol (Fig. 2C; Fig. 3).

**Figure 3.**
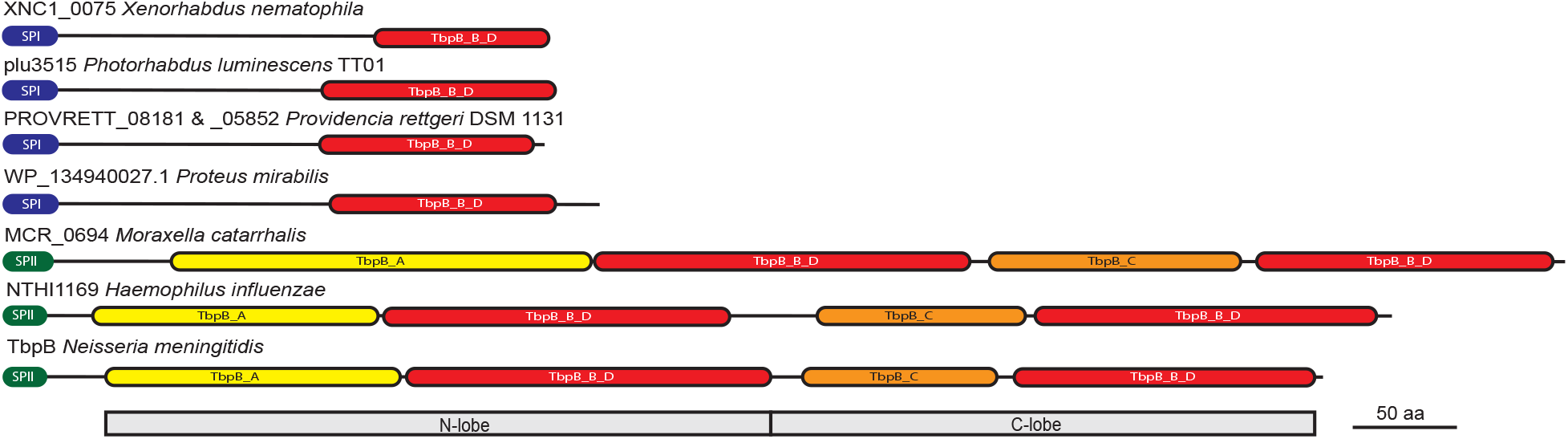
Examples of TbpBBDsol and TbpBBDlip domain architectures. The schematic diagram shows features found in select TbpBBD-domain containing proteins predicted to be exported by TXISS mechanisms. TXISS-1B associated proteins from *X. nematophila, P. luminescens, P. rettgeri, P. mirabilis*, and *H. haemolyticus* have signal-peptidase 1 (SPI) signal sequences, lack annotated “handle” domains, and have a single TbpBBD domain. TXISS-1A associated proteins from *M. catarrhalis, H. influenzae, and N. meningitidis* have signal-peptidase 2 (SPII) type signal sequences, N-lobe and C-lobe handle domains (TbpB_A and TbpB_C, respectively), and two TbpBBD domains. Predicted signal peptidase sites and domains are color-coded.

Biochemical and structural evidence support the conclusion that hemophilin is a soluble secreted protein that binds free heme and facilitates heme uptake into the cell (23). Three-dimensional homology modeling (Phyre^2^) was used to visualize potential structural similarities between hemophilin and several TbpBBDsol proteins (24, 25) (Supplemental Fig. 2). In light of sequence and structural level similarities we hypothesized that subcluster 1B proteins transport

TbpBBDsol proteins across the outer membrane via a mechanism analogous to subcluster 1A proteins transporting TbpBBDlip substrates. Our prediction separates the TXISS into two subfamilies, TXISS-1A which has members that flip TbpBBDlip substrates across the outer membrane and TXISS-1B which we predict has members that secrete TbpBBDsol substrates into the extracellular milieu.

### TXISS-1B activity reconstructed *in vivo*

To test our prediction that the TXISS-1B can secrete TbpBBDsol substrates, we investigated the <u>h</u>eme <u>r</u>eceptor <u>p</u>rotein (Hrp) locus of *X. nematophila*. This locus, which is conserved across the *Xenorhabdus* genus, consists of genes predicted to encode TonB, a TonB-dependent heme receptor named HrpA (XNC1_0073), the TXISS-1B homolog HrpB (XNC1_0074), and its predicted TbpBBDsol substrate HrpC (XNC1_0075) (11), a homolog of the heme-binding protein hemophilin (55% similar 39% identical) (23). Specifically, we sought to test whether the TXISS-1B homolog HrpB mediates secretion of the putative heme-binding protein HrpC. Using the expression vector pETDuet-1, HrpC-FLAG was expressed with and without HrpB-FLAG and whole cell and supernatant fractions were separated and analyzed by immunoblotting with anti-FLAG antibodies (26). We found that in the presence of HrpB-FLAG the levels of HrpC-FLAG detected in the supernatant increased 9.9-fold at 1-h post-induction and 17.0-fold at 2.5-h post-induction (Fig. 4A). Whole cell lysates demonstrated equivalent expression of HrpC-FLAG in both treatments (Fig. 4B). Coomassie staining of supernatants show that expression of HrpB-FLAG did not induce cell lysis or non-specific protein secretion (Fig. 4C, Supplemental Fig. 3). Consistent with the fact that in the absence of Slam1, several strains of *E. coli* surface expose a fraction of the substrate fHbp (27, 28), some HrpC-FLAG reached the supernatant in the absence of HrpB-FLAG.

**Figure 4.**
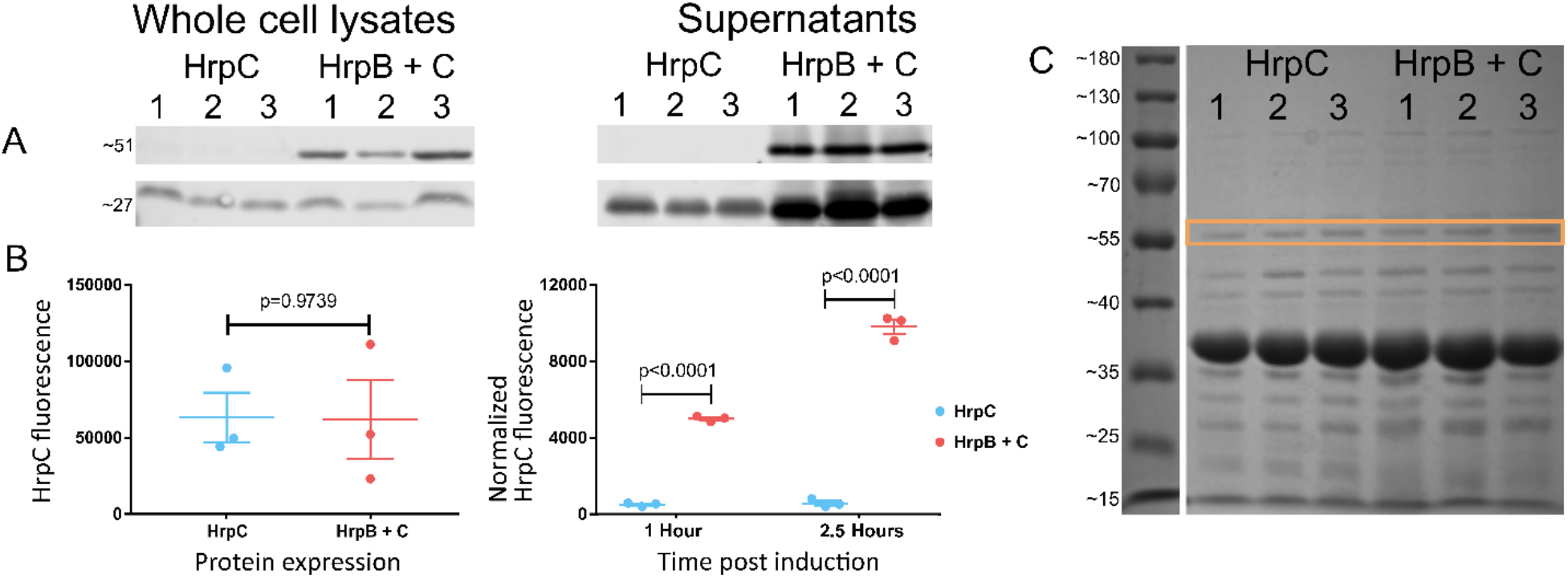
HrpB increases secretion of HrpC. A) Representative immunoblots of 2.5 h post-induction whole cell lysates and supernatant samples. HrpC-FLAG and HrpB-FLAG were distinguished by molecular weights (∼27 and ∼51 kDa, respectively). B) Fluorescence intensity of HrpC levels was not significantly impacted by HrpB in cell lysates (left graph, unpaired t-test) but was in supernatants (right graph, Sidak’s multiple comparisons test). C) Representative Coomassie stained SDS-PAGE gel of supernatant samples indicating HrpB causes neither cell lysis nor nonspecific protein export. A distinct band (in orange) was used as a loading control to normalize supernatant samples. The increase in supernatant HrpC is faintly visible at ∼27 kilodaltons in the last three supernatant lanes. Relicates labeled 1-3.

The Hrp locus is conserved in many proteobacteria including *Proteus mirabilis, Providencia rettgeri, Acinetobacter baumannii, N. gonorrhoeae*, and *H. haemolyticus*. For example, the TbpBBDsol protein present in *N. gonorrhoeae* (NGO0554) is flanked by a gene predicted to encode a TonB-dependent receptor (NGO0553) and a gene predicted to encode a DUF560 family protein (NGO0555). Consistent with the predicted role of these TbpBBDsol proteins in metal homeostasis, NGO0554 is repressed by iron, upregulated in response to oxidative stress, and contributes to resistance to peroxide challenge (29-31). These TXISS-dependent HrpC homologs likely contribute to metal homeostasis in a variety of human microbiome constituents by acting as hemophores akin to hemophilin and HasA (23, 32, 33).

### Host environment drives TXISS class

Having established that DUF560 homologs represent a bona fide secretion system, we next used the *Xenorhabdus* system to expand on our observation that the presence and type of TXISS corresponds to bacterial environmental niche. *Xenorhabdus* are species-specific obligate mutualists of *Steinernema* nematodes, and NilB is a known host range determinant. Therefore, we considered host species as an environmental niche and bioinformatically examined whether the complement of DUF560 genes in a microbe corresponds with host phylogeny (34-38). All 46 *Xenorhabdus* genomes on the Magnifying Genome (MaGe) platform (39) encode between one and three TXISS, with five distinct homologs represented across the genus (one TXISS-1A and four TXISS-1B) (40). Each unique combination of TXISS homologs was assigned one of six classes (A-F) as depicted in Supplemental Data Fig. 4 and Supplemental Table 3. To visualize correlations between TXISS class and host identity, we constructed Maximum Likelihood and Bayesian phylogenetic trees for *Xenorhabdus* and *Steinernema. Xenorhabdus* trees were generated with whole genome data while *Steinernema* trees used five loci as available (Supplemental Fig. 5; Supplemental Table 4). Aligning the Maximum likelihood phylogenies reveals that the TXISS complement of a *Xenorhabdus* species is more predictive of nematode host than phylogenetic position (Fig. 5).

**Figure 5.**
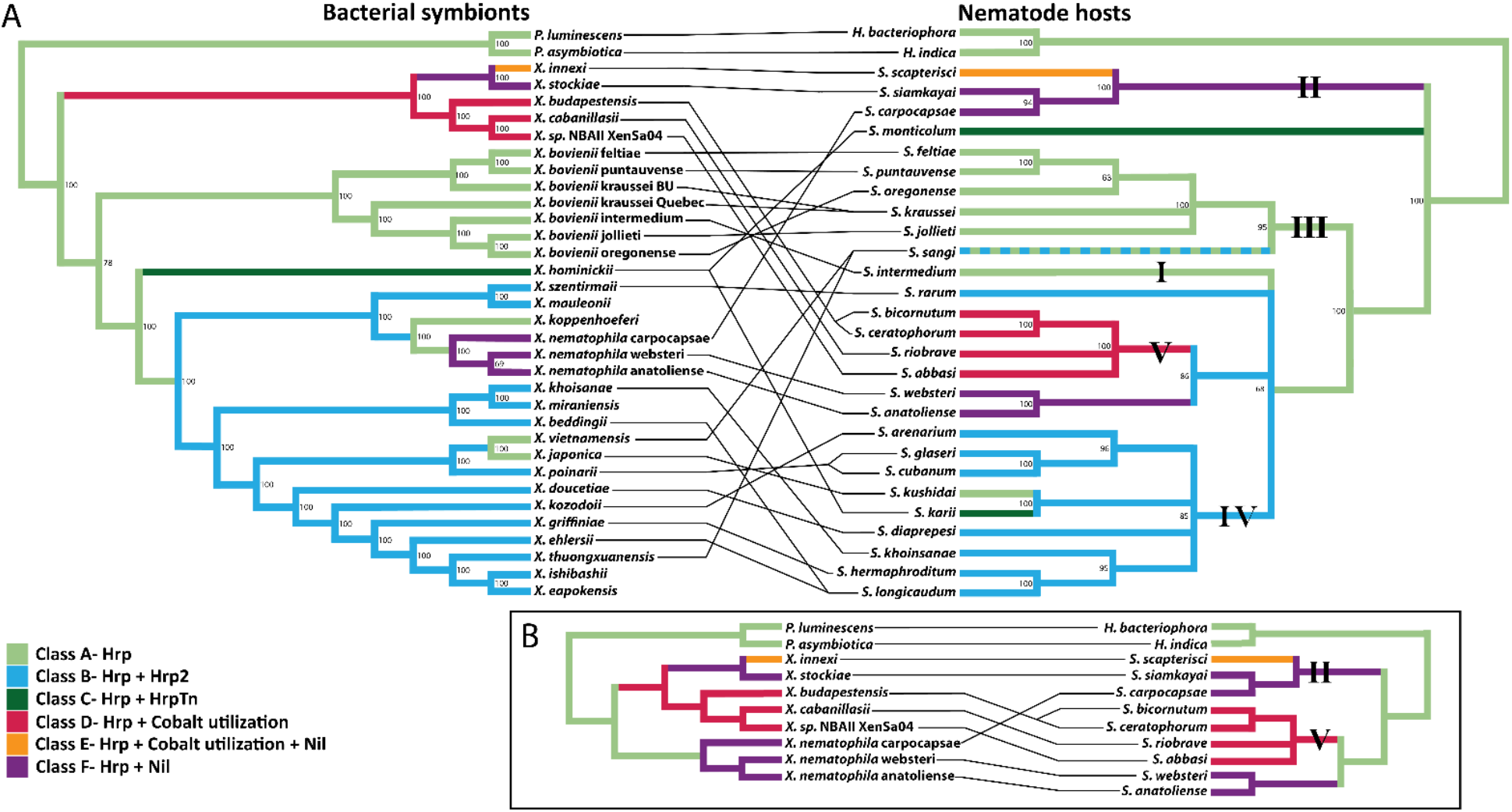
Cladograms of *Xenorhabdus* and *Steinernema* color coded according to *Xenorhabdus* DUF560 Class. Co-phylogeny of nematode species and their colonizing bacteria A) across the *Steinernema* genus or B) with a focus on specific clades. Numbers on branches indicate bootstrap support values. Bootstrap values below 60% were contracted. Lines connecting the phylogenies indicate mutualist pairs. Roman numerals highlight the 5 Steinernema clades described by Spridinov et al. 2004. Colored overlays indicate the DUF560 class of a given bacterium or a given nematode’s symbiont. Class A (light green); class B (light blue); class C (dark green); class D (red); class E (orange); class F (purple).

This alignment of the phylogenetic placement of a given *Steinernema* host with the TXISS complement of the symbiont provides insights into the nematode internal environment experienced by the symbiont. For example, most class B/C *Xenorhabdus* with two HrpB paralogs at the Hrp locus are symbionts of nematodes within the phylogenetic Clade IV, suggesting that these nematodes present a distinctive environment in which an additional Hrp locus is beneficial. Class D *Xenorhabdus* with an HrpB paralog encoded adjacent to genes predicted to encode a cobalt ABC transporter, are solely symbionts of Clade V nematodes. *X. innexi* and *X. stockiae* have seemingly diverged from this lineage through acquisition of a *nilB* homolog and switching to hosts within Clade II. Similarly, *X. nematophila* independently gained *nilB* and switched into a Clade II host (Fig. 5B). These acquisitions, alongside the varied genomic contexts of *nilB*/*nilC* pairs, are consistent with previous suggestions that the Nil locus was horizontally acquired among *Xenorhabdus* (14).

The TXISS NilB and the lipoprotein NilC are necessary for *X. nematophila* to colonize the Clade II nematode *S. carpocapsae* (10, 14). However, *X. nematophila* also colonizes two nematodes that are phylogenetically separate from Clade II, *S. anatoliense* and *S. websteri* (Fig. 5). Our hypothesis that TXISS are involved in host-environment adaptations leads to the prediction that *X. nematophila* will require the Nil locus to colonize these nematodes. To test this hypothesis, bacteria-free *S. anatoliense, S. websteri*, and *S. carpocapsae* eggs (41) were exposed to an *X. nematophila* ATCC19061 *Δnil* mutant and a Nil complemented strain (11). Consistent with our prediction, the *Δnil* mutant was deficient in infective juvenile colonization in all three nematode species (Fig. 6), demonstrating that *nil* genes are necessary for infective juvenile colonization of *S. anatoliense* and *S. websteri* and supporting our hypothesis that TXISS promote adaptation to host environments.

**Figure 6.**
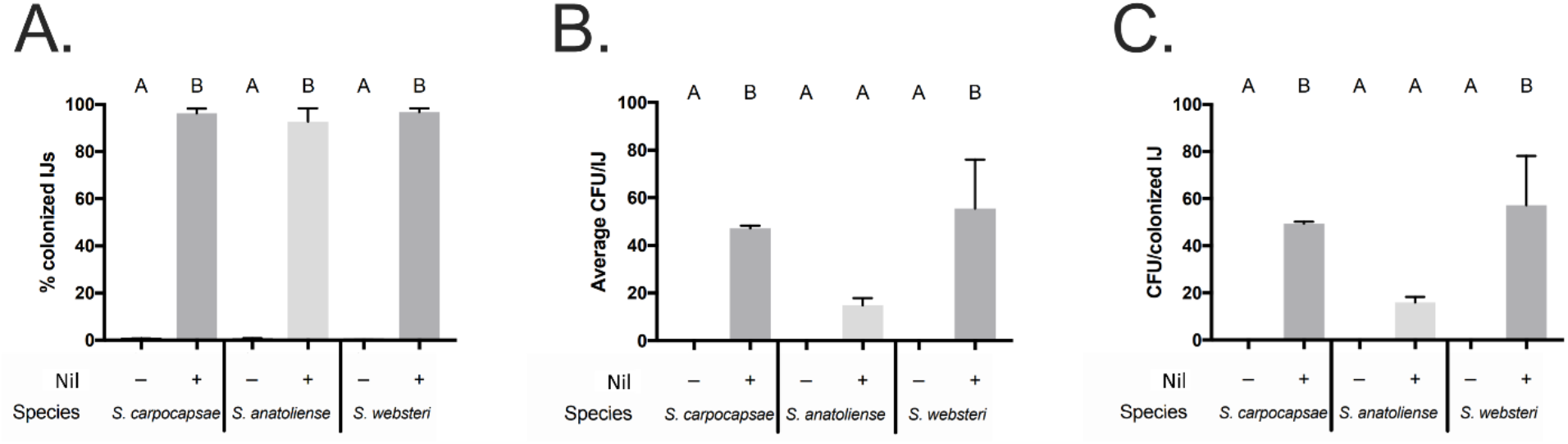
The Nil locus is necessary for colonization of *S. anatoliense, S. carpocapsae*, and *S. websteri* infective juveniles. Bacteria free *S. carpocapsae, S. anatoliense*, and *S. websteri* were exposed to GFP expressing *X. nematophila* lacking or bearing the Nil locus. The resulting progeny infective juveniles (IJs) were monitored for colonization either by A) microscopy or by B) plating lysates for average CFU/IJ. C) The average CFU per colonized IJ combines both of these values to show bacterial load per organism. Treatments were analyzed via one-way ANOVA and Tukey’s post-hoc test.

## Discussion

Bacteria that live inside animal hosts face many challenges as a result of their environment. Gram-negative bacteria that occupy host environments produce diverse factors that manipulate host cell and tissue physiology for their own benefit. Delivery of these effectors to their site of action requires the crossing of two bacterial membranes and possibly additional host cell membranes depending on the effector. Bacteria have evolved specialized secretion systems for delivery of effectors that facilitate the host-associated lifestyle. For instance, the type II secretion system can secrete hydrolyzing enzymes essential for commensals to grow, but it can also secrete clinically important toxins such as Cholera toxin and Diphtheria toxin (42, 43). Other secretion systems such as type III and type VI are specialized for the delivery of effectors across both bacterial and host cell membranes. Knowledge of cargo protein identities and sorting processes facilitates predictions from genomic information of bacterial secretome composition, regulation, and localization. Despite the diverse secretion systems now recognized, there are more to be identified based on the fact that some proteins predicted to be secreted lack known secretion pathways (44). The type X secretory pathway was described in 2020 and shed light on long standing mysteries surrounding the dependence on a muramidase for secretion of Typhoid toxin across the cell wall (45). Secretion pathways are the tools with which bacteria interact with their world, and as such each one discovered provides a slightly more complete picture of the toolbox.

The DUF560 (<u>d</u>omain of <u>u</u>nknown function) family presence in animal-associated bacteria was first recognized when it was noted that the host-colonization factor NilB has homologs in *N. meningitidis, M. catarrhalis, P. multocida*, and *H. influenzae* (10). This observation was strengthened by subsequent demonstration that Slam1 from human pathogenic bacteria facilitates the surface presentation of host-metal acquisition proteins (1, 2, 10). Enabled by the availability of extensive genomic data from many environments and ever-improving bioinformatic visualization tools, we have presented data demonstrating that the type XI secretion system is a broadly distributed molecular vehicle for moving proteins across the Gram-negative outer membrane. While DUF560 proteins were originally thought to represent a mechanism for lipoprotein surface exposure (1), recent studies have expanded that functional range to include peripheral membrane proteins (7) and now soluble secreted proteins, establishing the function of DUF560 OM proteins in secretion of varied substrates. We predict that further study will uncover a great diversity of cargo for the distinct classes of TXISS revealed through our network analysis. Here we focused on one single cluster of a DUF560 sequence similarity network and highlighted two subfamilies of TXISS (TXISS-1A and 1B). The remaining 9 clusters likely represent diverse subfamilies responsible for transporting as-yet-unknown cargo. The lipoprotein cargo proteins for which structural data are known revealed a common C-terminal 8-stranded β-barrel motif (2). Our discovery that HrpC is a cargo protein for TXISS-1B HrpB strengthens the concept that this barrel is an important characteristic of TXISS cargo; HrpC is a homolog of the *H. haemolyticus* hemophilin, the structure of which likewise adopts a C-terminal 8-stranded β-barrel (23) (Supplemental Fig. 2). These data support the concept that TXISS cargo have bifunctional structures in which the N-terminus is the host effector domain while the C-terminus directs the protein for secretion. This framework will facilitate identification of as-yet-unknown TXISS cargo among the genes that co-occur with TXISS OMPs. Further, our analyses suggest that sequence similarity network clusters have predictive power for other characteristics of TXISS cargo, including whether they are surface-attached or secreted. The network-enabled classification presented here will facilitate the investigation of both TXISS OMPs and their cargo in diverse bacteria.

Our work enables categorization of TXISS within *Neisseria* species that are found in many animal hosts, including the human pathogens *N. meningitidis* and *N. gonorrhoeae* (46). *Neisseria* strains can encode up to 6 TXISS paralogs. *N. meningitidis* MC58 has two functionally characterized TXISS: Slam1 and Slam2. Our network analysis indicates *N. meningitidis* encodes a third SPAM, NMB1466/ NP_274965, that also falls within Cluster 1A. *N. gonorrhoeae* TXISS are represented in more nodes than *N. meningitidis*, notably occupying 19.3% of nodes in subcluster 1B (Fig. 2). Taken together, these data indicate that *Neisseria* may be using TXISS to secrete both lipoproteins and soluble products.

There is a broadly conserved role for the TXISS in host-microbe interactions. Using the *Steinernema-Xenorhabdus* symbiosis we have demonstrated that the composition of TXISS in a bacterial symbiont genome correlates with host organism. During the evolutionary history of *Xenorhabdus* the gain or loss of TXISS loci correlated with host switching events (Fig. 5). Given the conservation of the TXISS Hrp locus, amongst all *Xenorhabdus* and throughout human microbiome constituents, it will be important in the future to examine the regulation of Hrp TXISS-dependent secretion and the roles of Hrp machinery in binding and acquiring host metals in a mucosal environment.

Finally, the whole family network first generated from the PF04575 (DUF560) dataset included cluster 5, comprising 14 nodes of *Klebsiella* homologs and 2 nodes with *Klebsiella* and *Escherichia* homologs. However, based on its limited number of nodes and predicted topology differences relative the rest of the network (16-stranded vs. 14-stranded barrel), cluster 5 was filtered out prior to our in-depth analysis. Cluster 5 homologs are annotated as PgaA/HmsH OMPs responsible for the synthase-dependent secretion of the exopolysaccharide poly-beta-1,6 N-acetyl-D-glucosamine (PNAG), which is necessary for biofilm formation (47, 48). The structure of the C-terminus of *E. coli* PgaA has been solved and forms a 16-stranded porin through which PNAG is exported (49). Although PgaA homologs exist in many bacteria, including *X. nematophila*, the only PgaA homologs represented in any cluster of the original network output were those from *Klebsiella* spp., further strengthening exclusion of cluster 5 from the TXISS network presented here (48). However, despite the topological and substrate (polysaccharide vs. protein) differences, the PF04575 assignment of cluster 5 members hints that there could be evolutionary or structural parallels between the PNAG exopolysaccharide (EPS) synthase-dependent secretion system and TXISS. PgaA is a component of one of several machineries responsible for export of EPS polymers including PNAG, colonic acid, alginate, cellulose, Psl, and Pel that comprise biofilm matrices of Gram-negative bacteria (47). In each case, like the TXISS system described here, the beta-barrel porin either has or associates with another protein that has TPR domains (47). The TPR region of *E. coli* PgaA is necessary for PNAG secretion and biofilm formation (49) and the TPR region of PelB modulates the activity of the polymer-modifying enzyme PelA, with impacts the chemistry of the resulting polymer (50). TXISS TPR domains may similarly modulate the activities of other proteins that influence secretion of the TXISS protein substrates. Indeed, *X. nematophila* expressing versions of the NilB TPR domain with small deletions display colonization defects that are ameliorated by deletion of the entire N-terminal periplasmic domain (11). It will be interesting for future studies to examine the commonalities between the TXISS and EPS synthase-dependent secretion systems and if there is any coordination in secretion of their respective substrates and/or host-associated phenotypes including aggregation and biofilm formation.

## Materials and Methods

### DUF560 sequence similarity network analysis

<u>E</u>nzyme <u>F</u>unction Initiative’s <u>E</u>nzyme <u>S</u>imilarity <u>T</u>ool (EFI-EST) was used to collect all predicted DUF560-domain-containing protein sequences from Interpro 73 and Uniprot 2019-02 (accessed 4.24.19) and BLAST for similarity (16, 17, 20). Representative networks collapsed nodes which shared ≥ 40% identity. On an EFI-EST network, edges are drawn according to a database-independent value called alignment score. A greater alignment score requirement means draws fewer edges. For separation of DUF560-domain-containing proteins an alignment score of 38 was chosen (Fig. 1). For subcluster 1A 89 was chosen and for subcluster 1B 100 was chosen (Supplemental Fig. 1). The EFI-EST Color SSN tool was used to assign cluster numbers. Networks were visualized and interpreted using Cytoscape v3.7.1 (51) and Gephi v0.9.2 (52). Nodes were arranged with the Fruchterman-Reingold force-directed layout algorithm (53).

The contents of each cluster were compared to Pfam DUF560 (PF04575) to ensure that clusters were legitimate DUF560 proteins (21). Any clusters for which fewer than 18% of sequences were present in Pfam, or which included fewer than 20 sequences were excluded from downstream analysis. This filtering removed cluster 5, which was composed mostly of *Klebsiella pneumoniae* PgaA. This generated a sequence similarity network that contains 10 clusters, 1222 nodes, and 52190 edges with 1589 TaxIDs represented (Fig. 1). Using NCBI Taxonomy Browser, each node was examined and manually curated for the isolates’ environmental origin(s) among the following categories: water, soil, plant, mammal, animal, invertebrate, nematode, built (environments such as sewage, bioreactors, etc.), multiple environments, and unclassified (Supplemental Table 1). If no citation was available, the isolation source was searched for in the biosamples or bioprojects records. If neither resource was available, strain source was searched for through other resources (NCBI linkout, Google search). A node was assigned an environment if the majority of strains within the node fell into that environment. If a node had no majority environment, the category “multiple environments” was assigned. Any node with animal associated microbes that did qualify as mammal, insect, or nematode associated was designated as generic “animal associated”. For fine scale interpretation, analysis focused on cluster 1 and its subclusters (Fig. 2; Supplemental Fig. 1)

Three different techniques were used to determine if DUF560 proteins present within our network were genomically associated with TbpB_B_D-domain-containing proteins, hereafter TbpBBD-domain-containing proteins. First, using EFI-GNN, genome-neighborhood-networks were generated for subcluster 1A (Alignment Score 89; 20 ORFs around), subcluster 1B (Alignment Score 38; 10 ORFs around), and subcluster 1C (Alignment Score 38; 10 ORFs around) resulting in 352 DUF560-TbpBBD pairs (17). Next, the RODEO web tool was used to analyze co-occurrence using profile Hidden Markov Models to assign domains to local ORFs resulting in 712 DUF560-TbpBBD pairs (22). 7 additional DUF560-TbpBBD pairs in *Xenorhabdus* were manually annotated. All three datasets were combined for a total of 851 non-redundant protein pairs (Supplemental Table 2). SignalP-5 (54, 55) was used to predict the signal peptides of all TbpBBD-domain-containing proteins. These predictions were used to annotate each node (Fig. 2C). Any node that was associated with both signal peptide bearing proteins and those with no predicted signal peptide were annotated according to the signal peptide bearing proteins.

### DUF560 genome neighborhood analysis

Subclusters 1A-C were separated and analyzed in EFI-EST with an alignment score of 38 as described above. Each network was then analyzed through EFI-GNN and visualized in Cytoscape v3.7.1 (51) (Alignment Score 38; 10 ORFs up and downstream).

### Phyre^2^ analysis

TbpBBDsol protein sequences from *Xenorhabdus nematophila* (HrpC), *Providencia rettgeri* (PROVETT_08181/PROVETT_05852), and *Proteus mirabilis* (WP_134940027.1) were collected and the first 22 amino acids were trimmed to remove the signal sequences. These sequences were entered into the Phyre^2^ Protein Homology/analogy Recognition Engine v.2.0 to predict potential 3-dimensional structures (25). The top predicted structural model output for all three proteins, hemophilin (6OM5), was used to generate structural models (Supplemental Fig. 2). Subsequent PDB files were visualized with Protean 3D v15. (Protean 3D®. Version 15.0. DNASTAR. Madison, WI).

### Co-expression of HrpB and HrpC

Primers and strains in Supplemental Table 5. Gene synthesis was performed by Genscript^®^ to synthesize the gene encoding HrpB25_26insDYKDDDDK (henceforth HrpB-FLAG). Genscript^®^ subsequently cloned HrpB-FLAG into multiple cloning site 2 (MCS2) of pETDuet-1. The genomic region containing HrpC was amplified from the *X. nematophila* ATCC 19061 (HGB800) genome using primers #11-12, digested with restriction enzymes SacI and SalI, and ligated into the multicloning site of pUC19. Site directed mutagenesis was used to add a C-terminal FLAG-tag onto HrpC using primers #13-14. The gene encoding HrpC1_2insV246_247insDYKDDDDK (henceforth HrpC-FLAG) was amplified from pUC19 using the primers #15-16. The gene product was ligated into multiple cloning site 1 (MCS1) of both pETDuet-1 and pETDuet-1/HrpB-FLAG, resulting in the expression plasmids pETDuet-1/HrpC-FLAG and pETDuet-1/HrpC-FLAG/HrpB-FLAG. Correct insertion of all genes was confirmed using primers #17-20 and Sanger sequencing at the University of Tennessee Genomics Core.

Expression plasmids were transformed into *E. coli* B21DE3 via electroporation (HGB2402 and HGB2459). All strains were grown solely in defined medium with 150 ug/ml ampicillin as previously described (56). Bacteria producing each expression plasmid were subcultured into 100 ml of broth at an initial OD600 of 0.028. Cultures were grown for 18 hours at 37°C to reach late logarithmic growth and then induced with 500 μM IPTG. One hour after induction a 700 μl sub-sample of each culture was taken and filter sterilized for subsequent protein precipitation. At 2.5 hours the whole cells were separated from the remaining supernatant and lysed open via a bead beater. Remaining supernatants were filter sterilized. Whole cell lysates were equalized via Bradford assay. For 1 and 2.5 hour supernatant samples 700 μl of supernatant was precipitated via 10% Trichloroacetic acid incubation (57).

Samples were run on 1% SDS-PAGE gels. For lysates, wells were loaded with 9.5ug of protein. For supernatants, wells were loaded with the precipitate of 350μl of media. Western blots were probed with a rat derived α-FLAG tag primary antibody and an α-IgG secondary antibody that fluoresces at 680 nm. Intensities for FLAG reactive bands were recorded. A distinct band from the Coomassie stained gel was used as a loading control to normalize intensities of supernatant samples prior to analysis (Fig. 4, Supplemental Fig. 3). For whole cell samples the data were analyzed with an unpaired t-test. For supernatant samples a two-way ANOVA was performed alongside Sidak’s multiple comparisons test.

### Phylogenetic tree generation

Phylogenetic analysis was performed as described previously (58). Briefly, select *Xenorhabdus* and *Photorhabdus* species were analyzed using MicroScope MaGe’s Gene Phyloprofile tool (34, 39) to identify homologous protein sets which were conserved across all assayed genomes. Putative paralogs were excluded from downstream analysis to ensure homolog relatedness, resulting in 665 homologous sets (Supplemental Table 5). Homolog sets were retrieved via locus tag indexing using BioPython (59), individually aligned using Muscle v3.8.31(60), concatenated using Sequence Matrix v1.8 (61), and trimmed of nucleotide gaps using TrimAL v1.3 (62). JmodelTest v2.1.10 (63) was used to choose the GTR+γ substitution model for maximum likelihood and Bayesian analysis.

For nematode phylogenetic analysis select *Steinernema* and *Heterorhabditis* species were analyzed. The Internal Transcribed Spacer, 18S rRNA, 28S rRNA, Cytochrome Oxidase I, and 12S rRNA loci were collected from Genbank’s as available and used as homologous sets (Supplemental Table 4). Nematode species which had fewer than 3 of the 5 loci sequenced were excluded from downstream analysis. Homologous sets were individually aligned, concatenated, and trimmed using the same methods as the *Xenorhabdus* sequences. JmodelTest v2.1.10 (63) was used to choose the GTR+γ+I substitution model.

Maximum likelihood analyses were performed via RAxML v8.2.10 (64) using rapid bootstrapping and 1000 replicates, and were visualized via Dendroscope v3.6.2 (65). Nodes with less than 60% bootstrap support were collapsed. Bayesian analyses were performed via MrBayes v3.2.6 with BEAGLE (66-68) on the Cipres Science Gateway platform (69). 500,000 MCMC replicates were performed for the bacterial tree, 4,000,000 were performed for the nematode tree. 25% of replicates were discarded as burn-in, and posterior probabilities were sampled every 500 replicates. Two runs were performed with 3 heated and one cold chain. The final standard deviation of split frequencies was 0.000000 for the bacterial tree, and 0.002557 for the nematode tree. Bayesian trees were visualized with FigTree v1.4.4 (70). Bayesian and maximum likelihood methods generated phylogenies with consistent topologies (Fig. 5; Supplemental Fig. 5).

### WT vs. *Δnil* colonization of *S. anatoliense, S. carpocapsae*, and *S. websteri*

Strains are described in Supplemental Table 5. Bacteria free eggs of *S. anatoliense, S. carpocapsae*, and *S. websteri* were generated (41) and exposed to a Nil locus mutant (HGB1495), and a Nil locus complemented strain (HGB1496) grown on lipid agar plates for 2 days at 25°C (11). Lipid agar plates were placed into White traps 1 week after plating eggs to collect infective juvenile (IJ) nematodes. Nematode colonization was visualized using fluorescence microscopy on the Keyence BZX-700 to observe GFP expressing bacteria in the receptacle. This was done for each species in biological triplicate and technical duplicate. To determine the number of CFU per IJ, nematodes were surface sterilized, and ground for 2 min using a Fisher brand motorized tissue grinder (CAT# 12-1413-61) to homogenize the nematodes and release colonizing bacteria. Serial dilutions in PBS were performed and plated on LB agar, which were incubated at 30°C for 1 day before enumerating CFUs (Fig. 6). To calculate the CFU per colonized IJ, the percent colonized nematodes was divided by the CFU/IJ for each biological replicate. The data were analyzed using a one-way ANOVA with Tukey’s multiple comparison’s test to compare the mean of each treatment.

## Acknowledgements

This work was supported by a grant from the National Science Foundation (IOS-1353674) to HGB and KTF and the University of Tennessee-Knoxville to HGB. TJM was supported by an NIH National Research Service Award T32-GM07215.

## Author Contributions

TJM, ASG, and HGB wrote the text of this article and composed the figures. Bioinformatic analysis was performed by ASG and TJM. Phylogenetic analysis was performed by ASG. Cloning and experiments were performed by ASG and TJM. KTF provided sustained intellectual contributions to the research design and edits to the text and figures.

## Additional Information

The authors have no competing interests that might be perceived to influence the results and/or discussion reported in this paper. A previous version of this article was published on BioRχiv (https://doi.org/10.1101/2020.01.20.912956). Correspondence and requests for materials should be addressed to H. Goodrich-Blair at

## Supplemental Data

**Supplemental Figure 1.**
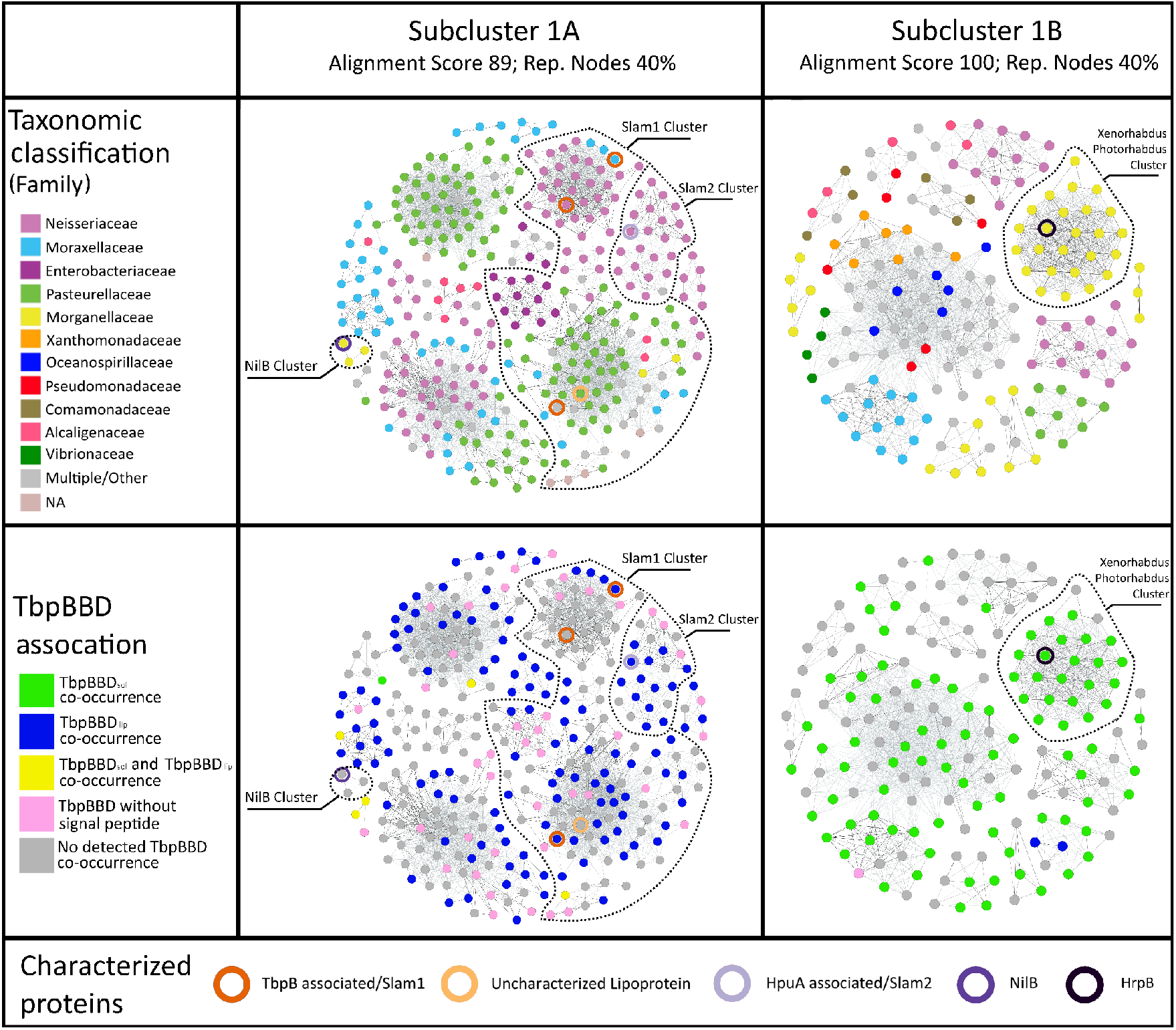
Increased stringency separates subcluster 1A and 1B into functional groups. A series of stringent EFI-EST sequence similarity networks highlights detail in cluster 1 of the DUF560 homologs. Edge darkness demonstrates similarity. Node positioning was optimized using the Fruchterman-Reingold algorithm (53). Dotted lines indicate hypothetical functional clusters based on previous molecular data. Circled nodes indicate proteins which have been molecularly characterized. Subclusters 1A and 1B were analyzed separately to allow fine tuning of alignment score (89 and 100 respectively). Networks were color coded to display either taxonomic categories or co-inheritance with TbpBBD-domain containing proteins.

**Supplemental Figure 2.**
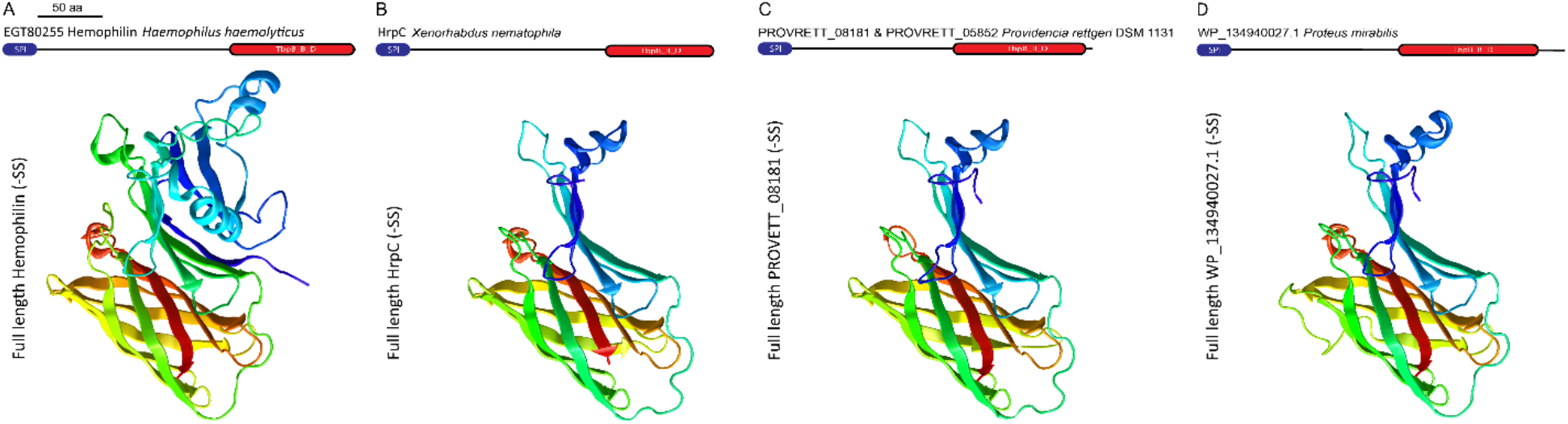
Phyre^2^ models of select TbpBBD_sol_ proteins. TbpBBD_sol_ proteins, lacking the signal sequence (-SS), from *X. nematophila* (HrpC), *P. rettgeri* (PROVRETT_08181 and 05852), and *P. mirabilis* (WP_134940027.1) were queried through the Phyre^2^ Protein Homology/analogy Recognition Engine v. 2.0 (http://www.sbg.bio.ic.ac.uk/phyre2/html/page.cgi?id=index) (25). The top predicted structural model output for each is shown alongside the solved crystal structure of hemophilin from *H. haemolyticus* (protein data bank file 6OM5) which the algorithm selected as the template for all queries (23).

**Supplemental Figure 3.**
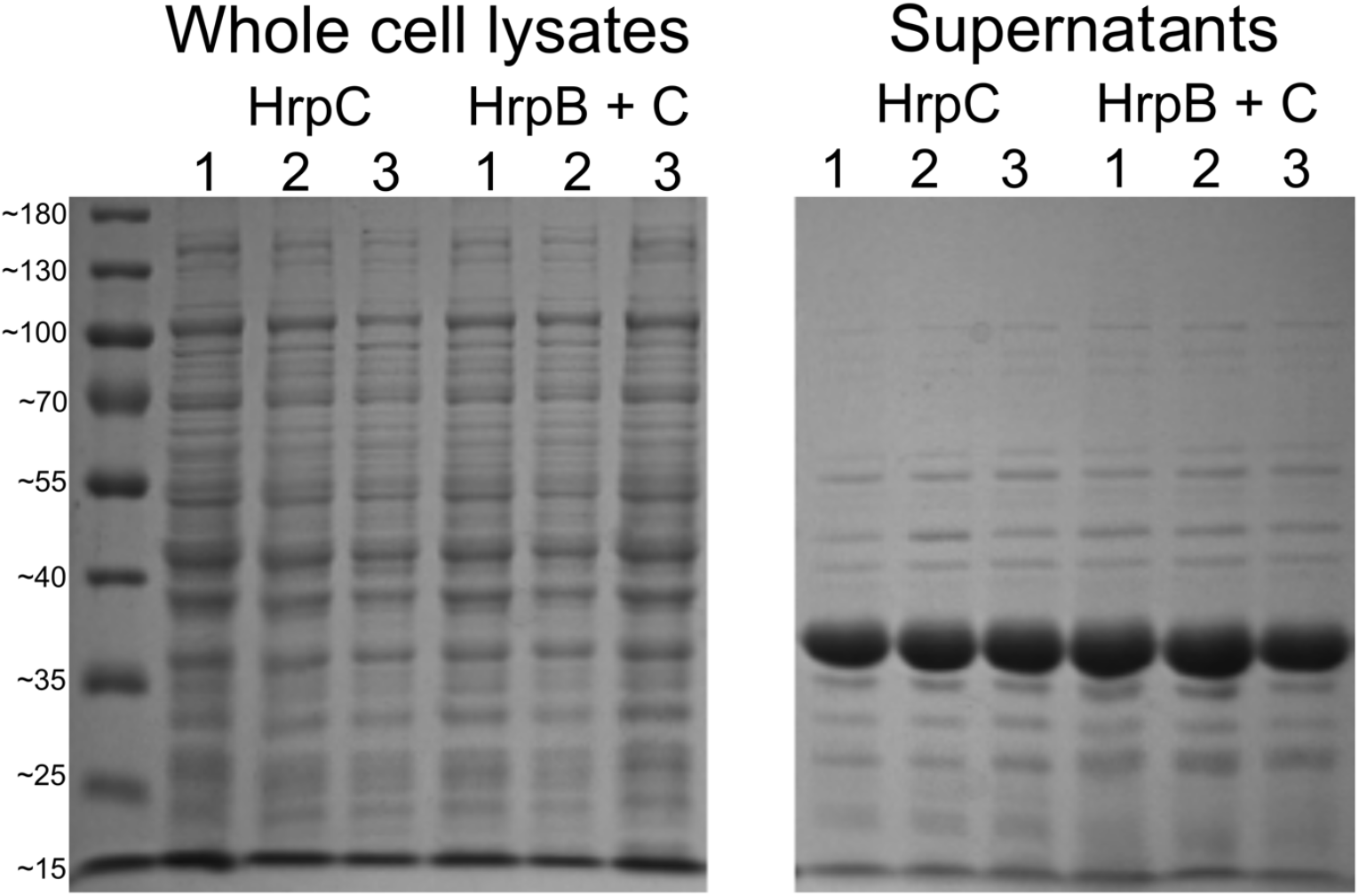
Supernatant samples have distinct protein profile. Coomassie stained SDS-PAGE gels showing side by side comparison of protein banding in whole cell lysates and supernatant samples of *E. coli* expressing the indicated proteins. Units on the left side are in kilodaltons. Expression of HrpB does not cause cell lysis or nonspecific protein export, if it did the banding pattern of the supernatant would look more like that of the lysates. The increase in supernatant HrpC is faintly visible at ∼27 kilodaltons in the last three supernatant sample lanes. All experiments were repeated in biological triplicate as labeled above the lanes.

**Supplemental Figure 4.**
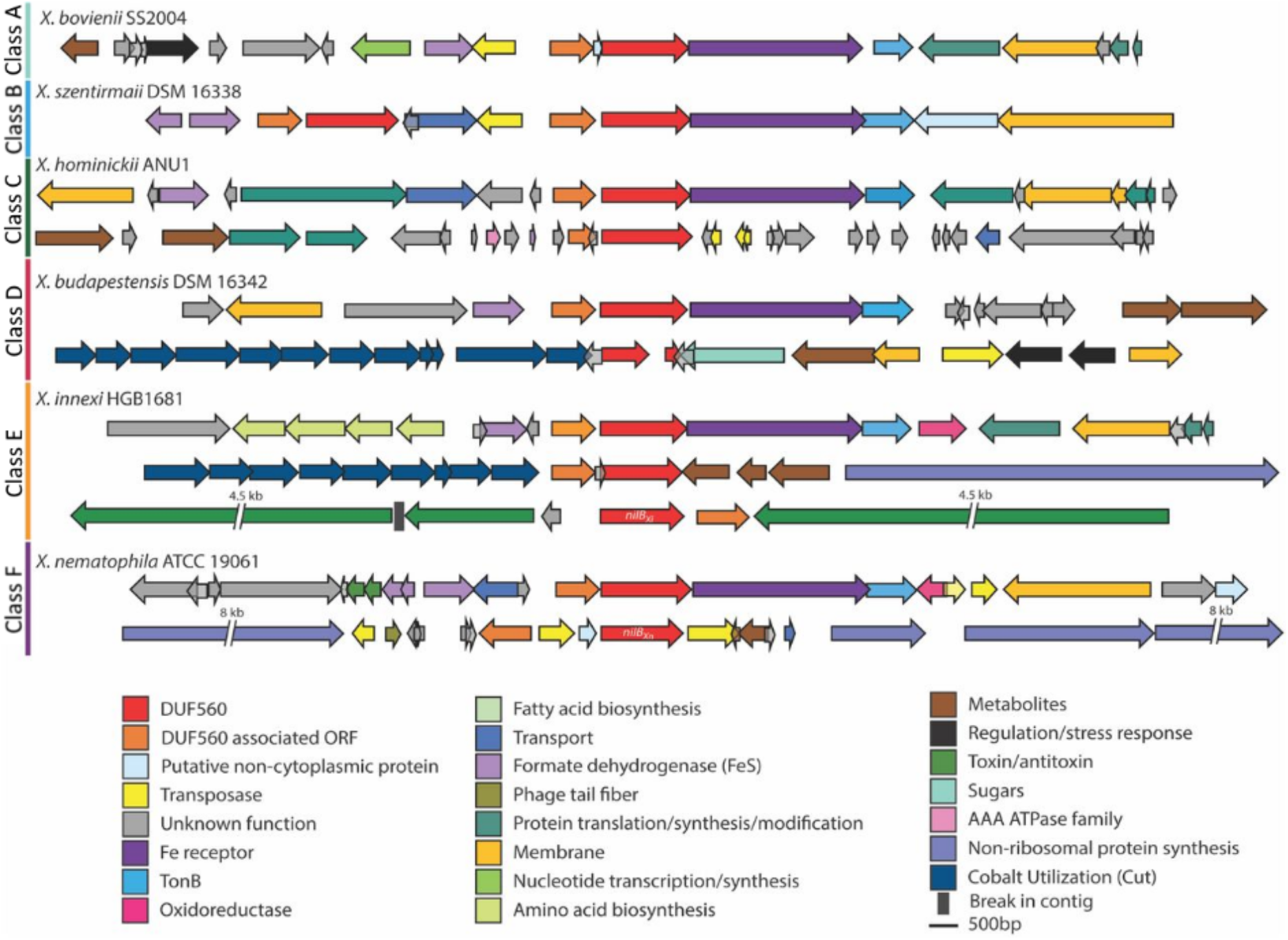
Representative genomes of *Xenorhabdus* DUF560 Classes A-F. Schematic diagrams of *Xenorhabdus* TXISS loci representing each of the six classes defined in the text. One species from each of the classes was selected for presentation. Box arrows represent open reading frames (ORFs), which are color coded according to predicted annotated function as indicated by the legend. The DUF560 homolog is shown in red and the predicted TXISS cargo is shown in orange. Large ORFs were not presented in their entirety and the length of the gap is indicated above the break line shown within such ORFs.

**Supplemental Figure 5.**
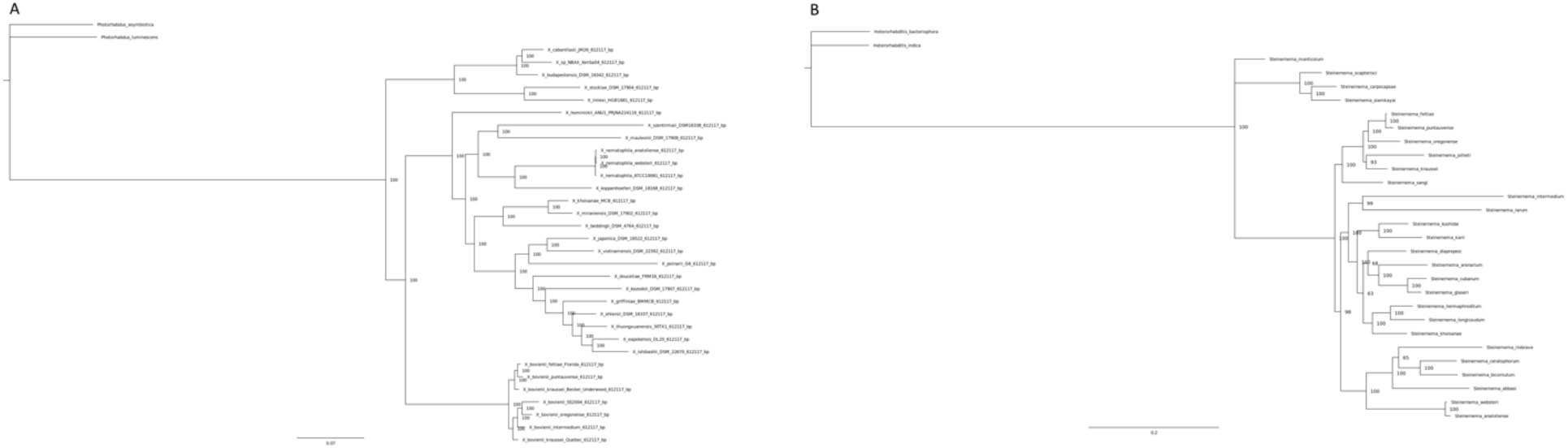
Bayesian posterior probability phylogenies. A) Phylogram of select *Xenorhabdus* bacteria, based on concatenations of 665 conserved core genes. Numbers indicate posterior probability values. Distances indicate substitutions per base pair. B) Bayesian phylogeny of select entomopathogenic nematodes, based on concatenations of the ITS, 18S rRNA, 28S rRNA, COI, and 12S rRNA loci. Two members of the sister taxon *Photorhabdus* were chosen as an outgroup. Loci are recorded in Supplemental table 4.

**Supplemental Table 1. DUF560 sequence similarity network clusters 1-4 and 6-11 node table, with subcluster, environmental annotations, and sources (when possible)**

*Associated file: TableS1_Clusters1-4,6-11WithEnvTax*.*xlsx*

**Supplemental Table 2. DUF560 TbpBBD protein Co-occurrence**

*Associated file: TableS2_DUF560TbpBBDCo-occurrence*.*xlsx*

**Supplemental Table 3. *Xenorhabdus* DUF560 classes**

*Associated file: TableS3_XenorhabdusDUF560Classes*.*xlsx*

**Supplemental Table 4. Loci tags used in phylogenetic analysis of *Xenorhabdus* and *Steinernema* species**

*Associated file: TableS4_CombinedLociTable*

**Supplemental Table 5. Primer and strain list**

*Associated file: TableS5_PrimersAndStrains*.*xlsx*

## References

1. Hooda Y, Lai CC, Judd A, Buckwalter CM, Shin HE, Gray-Owen SD, Moraes TF. 2016. Slam is an outer membrane protein that is required for the surface display of lipidated virulence factors in Neisseria. Nat Microbiol 1:16009.

2. Hooda Y, Lai CCL, Moraes TF. 2017. Identification of a Large Family of Slam-Dependent Surface Lipoproteins in Gram-Negative Bacteria. Front Cell Infect Microbiol 7:207.

3. Chu BC, Garcia-Herrero A, Johanson TH, Krewulak KD, Lau CK, Peacock RS, Slavinskaya Z, Vogel HJ. 2010. Siderophore uptake in bacteria and the battle for iron with the host; a bird’s eye view. Biometals 23:601–11.

4. Noinaj N, Guillier M, Barnard TJ, Buchanan SK. 2010. TonB-dependent transporters: regulation, structure, and function. Annual Review of Microbiology 64:43–60.

5. Schweppe DK, Harding C, Chavez JD, Wu X, Ramage E, Singh PK, Manoil C, Bruce JE. 2015. Host-Microbe Protein Interactions during Bacterial Infection. Chem Biol 22:1521–1530.

6. Chavez-Dozal AA, Gorman C, Lostroh CP, Nishiguchi MK. 2014. Gene-swapping mediates host specificity among symbiotic bacteria in a beneficial symbiosis. PLoS One 9:e101691.

7. da Silva RAG, Karlyshev AV, Oldfield NJ, Wooldridge KG, Bayliss CD, Ryan A, Griffin R. 2019. Variant Signal Peptides of Vaccine Antigen, FHbp, Impair Processing Affecting Surface Localization and Antibody-Mediated Killing in Most Meningococcal Isolates. Front Microbiol 10:2847.

8. Akhurst RJ. 1983. Neoplectana species: specificity of association with bacteria of the genus Xenorhabdus. Exp Parasitol 55:258–263.

9. Chaston JM, Suen G, Tucker SL, Andersen AW, Bhasin A, Bode E, Bode HB, Brachmann AO, Cowles CE, Cowles KN, Darby C, de Leon L, Drace K, Du Z, Givaudan A, Herbert Tran EE, Jewell KA, Knack JJ, Krasomil-Osterfeld KC, Kukor R, Lanois A, Latreille P, Leimgruber NK, Lipke CM, Liu R, Lu X, Martens EC, Marri PR, Medigue C, Menard ML, Miller NM, Morales-Soto N, Norton S, Ogier JC, Orchard SS, Park D, Park Y, Qurollo BA, Sugar DR, Richards GR, Rouy Z, Slominski B, Slominski K, Snyder H, Tjaden BC, van der Hoeven R, Welch RD, Wheeler C, Xiang B, Barbazuk B, et al. 2011. The entomopathogenic bacterial endosymbionts Xenorhabdus and Photorhabdus: convergent lifestyles from divergent genomes. PLoS One 6:e27909.

10. Heungens K, Cowles CE, Goodrich-Blair H. 2002. Identification of Xenorhabdus nematophila genes required for mutualistic colonization of Steinernema carpocapsae nematodes. Mol Microbiol 45:1337–53.

11. Bhasin A, Chaston JM, Goodrich-Blair H. 2012. Mutational analyses reveal overall topology and functional regions of NilB, a bacterial outer membrane protein required for host association in a model of animal-microbe mutualism. J Bacteriol 194:1763–76.

12. Blatch GL, Lassle M. 1999. The tetratricopeptide repeat: a structural motif mediating protein-protein interactions. Bioessays 21:932–9.

13. Cowles CE, Goodrich-Blair H. 2004. Characterization of a lipoprotein, NilC, required by Xenorhabdus nematophila for mutualism with its nematode host. Mol Microbiol 54:464–77.

14. Cowles CE, Goodrich-Blair H. 2008. The Xenorhabdus nematophila nilABC genes confer the ability of Xenorhabdus spp. to colonize Steinernema carpocapsae nematodes. J Bacteriol 190:4121–8.

15. d’Enfert C, Chapon C, Pugsley AP. 1987. Export and secretion of the lipoprotein pullulanase by Klebsiella pneumoniae. Mol Microbiol 1:107–16.

16. Gerlt JA, Bouvier, J.T., Davidson, D.B., Imker, H.J., Sadkhin, B., Slater, D.R., Whalen, K.L. 2015. Enzyme Function Initiative-Enzyme Similarity Tool (EFI-EST): A web tool for generating protein sequence similarity networks. Biochemica et Biophysica Acta 1854:1019–1037.

17. Zallot R, Oberg, N.O., Gerlt, J.A. 2018. ‘Democratized’genomicenzymologywebtoolsforfunctionalassignment. Current Opinion in Chemical Biology 47:77–85.

18. Zallot R, Oberg N, Gerlt JA. 2019. The EFI Web Resource for Genomic Enzymology Tools: Leveraging Protein, Genome, and Metagenome Databases to Discover Novel Enzymes and Metabolic Pathways. Biochemistry 58:4169–4182.

19. Ostberg KL, DeRocco, A.J., Mistry, S.D., Dickinson, M.K., Cornelissen, C.N. 2013. Conserved Regions of Gonococcal TbpB Are Critical for Surface Exposure and Transferrin Iron Utilization. Infection and Immunity 81:3442–3450.

20. Gerlt JA. 2017. Genomic Enzymology: Web Tools for Leveraging Protein Family Sequence−Function Space and Genome Context to Discover Novel Functions. Biochemistry 56:4293–4308.

21. El-Gebali S, Mistry J, Bateman A, Eddy SR, Luciani A, Potter SC, Qureshi M, Richardson LJ, Salazar GA, Smart A, Sonnhammer ELL, Hirsh L, Paladin L, Piovesan D, Tosatto SCE, Finn RD. 2019. The Pfam protein families database in 2019. Nucleic Acids Research 47:D427–D432.

22. Choe K, Mitchell Laboratory. 2018. RODEO (Rapid ORF Description & Evaluation Online) is an algorithm to help biosynthetic gene cluster (BGC) analysis, with an emphasis on ribosomal natural product (RiPP) discovery., p http://ripp.rodeo/advanced.html, University of Illinois at Urbana-Champaign.

23. Latham RD, Torrado M, Atto B, Walshe JL, Wilson R, Guss JM, Mackay JP, Tristram S, Gell DA. 2020. A heme-binding protein produced by Haemophilus haemolyticus inhibits non-typeable Haemophilus influenzae. Mol Microbiol 113:381–398.

24. Kelley LA, Sternberg MJ. 2009. Protein structure prediction on the Web: a case study using the Phyre server. Nat Protoc 4:363–71.

25. Kelley LA, Mezulis S, Yates CM, Wass MN, Sternberg MJ. 2015. The Phyre2 web portal for protein modeling, prediction and analysis. Nat Protoc 10:845–58.

26. Einhauer A, Jungbauer A. 2001. The FLAG peptide, a versatile fusion tag for the purification of recombinant proteins. J Biochem Biophys Methods 49:455–65.

27. Fantappie L, Irene C, De Santis M, Armini A, Gagliardi A, Tomasi M, Parri M, Cafardi V, Bonomi S, Ganfini L, Zerbini F, Zanella I, Carnemolla C, Bini L, Grandi A, Grandi G. 2017. Some Gram-negative lipoproteins keep their surface topology when transplanted from one species to another and deliver foreign polypeptides to the bacterial surface. Mol Cell Proteomics 16:1348–1364.

28. Konar M, Rossi R, Walter H, Pajon R, Beernink PT. 2015. A mutant library approach to identify improved meningococcal factor H binding protein vaccine antigens. PLoS One 10:e0128185.

29. Jackson LA, Ducey TF, Day MW, Zaitshik JB, Orvis J, Dyer DW. 2010. Transcriptional and functional analysis of the Neisseria gonorrhoeae Fur regulon. J Bacteriol 192:77–85.

30. Quillin SJ, Hockenberry AJ, Jewett MC, Seiferta HS. 2018. Neisseria gonorrhoeae exposed to sublethal levels of hydrogen peroxide mounts a complex transcriptional tesponse. mSystems 3:e00156–18.

31. Stohl EA, Criss AK, Seifert HS. 2005. The transcriptome response of Neisseria gonorrhoeae to hydrogen peroxide reveals genes with previously uncharacterized roles in oxidative damage protection. Mol Microbiol 58:520–532.

32. Dent AT, Mouriño S, Huang W, Wilks A. 2018. Post-transcriptional regulation of the Pseudomonas aeruginosa heme assimilation system (Has) fine-tunes extracellular heme sensing. J Biol Chem 294:2771–2785.

33. Yukl ET, Jepkorir G, Alontaga AY, Pautsch L, Rodriguez JC, Rivera M, Moënne-Loccoz P. 2010. Kinetic and spectroscopic studies of hemin acquisition in the hemophore HasAp from Pseudomonas aeruginosa. Biochemistry 49:6646–6654.

34. Chaston JM, Murfin KE, Heath-Heckman EA, Goodrich-Blair H. 2013. Previously unrecognized stages of species-specific colonization in the mutualism between Xenorhabdus bacteria and Steinernema nematodes. Cellular Microbiology 15:1545–1559.

35. Kampfer P, Tobias NJ, Ke LP, Bode HB, Glaeser SP. 2017. Xenorhabdus thuongxuanensis sp. nov. and Xenorhabdus eapokensis sp. nov., isolated from Steinernema species. International Journal of Systematic and Evolutionary Microbiology 67:1107–1114.

36. Lee MM, Stock SP. 2010. A multigene approach for assessing evolutionary relationships of Xenorhabdus spp. (gamma-Proteobacteria), the bacterial symbionts of entomopathogenic Steinernema nematodes. J Invertebr Pathol 104:67–74.

37. Snyder HA, Stock SP, Kim SK, Flores-Lara Y, Forst S. 2007. New insights into the colonization and release process of Xenorhabdus nematophila and the morphology and ultrastructure of the bacterial receptacle of its nematode host, Steinernema carpocapsae. Applied and Environmental Microbiology 73:5338–5346.

38. Spiridonov SE, Reid AP, Podrucka K, Subbotin SA, Moens M. 2004. Phylogenetic relationships within the genus Steinernema (Nematoda: Rhabditida) as inferred from analyses of sequences of the ITS1-5.8S-ITS2 region of rDNA and morphological features. Nematology 6:547–566.

39. Vallenet D, Calteau A, Cruveiller S, Gachet M, Lajus A, Josso A, Mercier J, Renaux A, Rollin J, Rouy Z, Roche D, Scarpelli C, Medigue C. 2017. MicroScope in 2017: an expanding and evolving integrated resource for community expertise of microbial genomes. Nucleic Acids Research 45:D517–D528.

40. McGinnis S, Madden TL. 2004. BLAST: at the core of a powerful and diverse set of sequence analysis tools. Nucleic Acids Res 32:W20–5.

41. Murfin KE, Chaston J, Goodrich-Blair H. 2012. Visualizing bacteria in nematodes using fluorescence microscopy. J Vis Exp 68:e4298.

42. Costa TR, Felisberto-Rodrigues C, Meir A, Prevost MS, Redzej A, Trokter M, Waksman G. 2015. Secretion systems in Gram-negative bacteria: structural and mechanistic insights. Nat Rev Microbiol 13:343–59.

43. Zielke RA, Simmons RS, Park BR, Nonogaki M, Emerson S, Sikora AE. 2014. The type II secretion pathway in Vibrio cholerae is characterized by growth phase-dependent expression of exoprotein genes and is positively regulated by sigmaE. Infect Immun 82:2788–801.

44. Bendtsen JD, Kiemer L, Fausboll A, Brunak S. 2005. Non-classical protein secretion in bacteria. BMC Microbiol 5:58.

45. Palmer T, Finney AJ, Saha CK, Atkinson GC, Sargent F. 2020. A holin/peptidoglycan hydrolase-dependent protein secretion system. Mol Microbiol doi:10.1111/mmi.14599.

46. Cohen J, Powderly W, Opal S. 2017. Preface to the Fourth Edition, Infectious Diseases, 4th edition. Elsevier.

47. Low KE, Howell PL. 2018. Gram-negative synthase-dependent exopolysaccharide biosynthetic machines. Curr Opin Struct Biol 53:32–44.

48. Whitney JC, Howell PL. 2013. Synthase-dependent exopolysaccharide secretion in Gram-negative bacteria. Trends Microbiol 21:63–72.

49. Wang Y, Andole Pannuri A, Ni D, Zhou H, Cao X, Lu X, Romeo T, Huang Y. 2016. Structural Basis for Translocation of a Biofilm-supporting Exopolysaccharide across the Bacterial Outer Membrane. J Biol Chem 291:10046–57.

50. Marmont LS, Whitfield GB, Rich JD, Yip P, Giesbrecht LB, Stremick CA, Whitney JC, Parsek MR, Harrison JJ, Howell PL. 2017. PelA and PelB proteins form a modification and secretion complex essential for Pel polysaccharide-dependent biofilm formation in Pseudomonas aeruginosa. J Biol Chem 292:19411–19422.

51. Shannon P, Markiel A, Ozier O, Baliga NS, Wang JT, Ramage D, Amin N, Schwikowski B, Ideker T. 2003. Cytoscape: a software environment for integrated models of biomolecular interaction networks. Genome Res 13:2498–504.

52. Bastian M, Heymann S, Jacomy M. 2009. Gephi: An Open Source Software for Exploring and Manipulating Networks. International AAAI Conference on Weblogs and Social Media.

53. Fruchterman TMJ, Reingold EM. 1991. Graph Drawing by Force-directed Placement. Software -Practice and Experience 21:1129–1164.

54. Almagro Armenteros JJ, Tsirigos KD, Sonderby CK, Petersen TN, Winther O, Brunak S, von Heijne G, Nielsen H. 2019. SignalP 5.0 improves signal peptide predictions using deep neural networks. Nat Biotechnol 37:420–423.

55. Nielsen H, Engelbrecht J, Brunak S, von Heijne G. 1997. Identification of prokaryotic and eukaryotic signal peptides and prediction of their cleavage sites. Protein Engineering 10:1–6.

56. Orchard SS, Goodrich-Blair H. 2004. Identification and functional characterization of a Xenorhabdus nematophila oligopeptide permease. Appl Environ Microbiol 70:5621–7.

57. Koontz L. 2014. TCA Precipitation, p 3–10. In Lorsch J (ed), Laboratory Methods in Enzymology: Protein Part C, vol 536. Academic Press Elsevier.

58. Murfin KE, Lee MM, Klassen JL, McDonald BR, Larget B, Forst S, Stock SP, Currie CR, Goodrich-Blair H. 2015. Xenorhabdus bovienii strain diversity impacts coevolution and symbiotic maintenance with Steinernema spp. nematode hosts. MBio 6:e00076.

59. Cock PJ, Antao T, Chang JT, Chapman BA, Cox CJ, Dalke A, Friedberg I, Hamelryck T, Kauff F, Wilczynski B, de Hoon MJ. 2009. Biopython: freely available Python tools for computational molecular biology and bioinformatics. Bioinformatics 25:1422–3.

60. Edgar RC. 2004. MUSCLE: multiple sequence alignment with high accuracy and high throughput. Nucleic Acids Res 32:1792–7.

61. Vaidya G, Lohman, D. J., Meier, R. 2010. SequenceMatrix: Concatenation software for the fast assembly of multi-gene datasets with character set and codon information. Cladistics 27:171–180.

62. Capella-Gutiérrez S, Silla-MartÍnez, J. M., Gabaldón, T. 2009. trimAl: a tool for automated alignment trimming in large-scale phylogenetic analyses. Bioinformatics 25:1972–1973.

63. Darriba D, Taboada, G. L., Doallo, R., Posada, D. 2012. jModelTest 2: more models, new heuristics and highperformance computing. Nat Methods 9:772.

64. Stamatakis A. 2014. RAxML version 8: a tool for phylogenetic analysis and post-analysis of large phylogenies. Bioinformatics 30:1312–1313.

65. Huson DH, Scornavacca, C. 2012. Dendroscope 3: An Interactive Tool for Rooted Phylogenetic Trees and Networks. Syst Biol 61:1061–1067.

66. Altekar G, Dwarkadas, S., Huelsenbeck, J. P., Ronquist, F. 2004. Parallel Metropolis coupled Markov chain Monte Carlo for Bayesian phylogenetic inference Bioinformatics 20:407–415.

67. Ayres DL, Darling, A., Zwickl, D. J., Beerli, P., Holder, M. T., Lewis, P. O., Huelsenbeck, J. O., Ronquist, F., Swofford, D. L., Cummings, M. P., Rambaut, A., Suchard, M. A. 2011. BEAGLE: An Application Programming Interface and High-Performance Computing Library for Statistical Phylogenetics. Syst Biol 61:170–173.

68. Ronquist F, Teslenko, M., van der Mark, P., Ayres, D. L., Darling, A., Hohna, S., Larget, B., Liu, L., Suchard, M. A., Huelsenbeck, J. P. 2012. MrBayes 3.2: Efficient Bayesian Phylogenetic Inference and Model Choice Across a Large Model Space. Syst Biol 61:539–542.

69. Miller MA, Pfeiffer, W., Schwartz, T. Creating the CIPRES Science Gateway for inference of large phylogenetic trees, p 1–8. In (ed), Proceedings of the Gateway Computing Environments Workshop (GCE),

70. Rambaut A. 2018. FigTree v1.4.4 p http://tree.bio.ed.ac.uk/software/figtree/.

